# Healing of ischemic injury in the retina

**DOI:** 10.1101/2024.11.04.621932

**Authors:** Silke Becker, Jordan Allen, Zia L’Ecuyer Morison, Sama Saeid, Austin Adderley, Ari Koskelainen, Frans Vinberg

## Abstract

Neuro- and retinal degenerative diseases, including Alzheimer’s, age-related macular degeneration, stroke, and central retinal artery occlusion, rob millions of their independence. Studying these diseases in human retinas has been hindered by the immediate loss of neuronal activity postmortem. While recent studies restored limited activity in postmortem CNS tissues, synchronized neuronal transmission >30 minutes postmortem remained elusive. Our study overcomes this barrier by reviving and sustaining light signal transmission in human retinas recovered up to four hours and stored 48 hours postmortem. We also establish infrared-based *ex vivo* imaging for precise sampling, a closed perfusion system for drug testing, and an *ex vivo* ischemia-reperfusion model in mouse and human retina. This platform enables testing of neuroprotective and neurotoxic effects of drugs targeting oxidative stress and glutamate excitotoxicity. Our advances question the irreversibility of ischemic injury, support preclinical vision restoration studies, offer new insights into treating ischemic CNS injuries, and pave the way for transplantation of human donor eyes.

**Teaser:** Reviving light signaling in postmortem human retinas challenges the irreversibility of ischemic injury and advances research to restore vision.

## INTRODUCTION

Until recently, consensus decreed that neurons in the central nervous system (CNS), including the retina, rapidly and irreversibly deteriorate after the blood circulation ceases [1], [2], [3]. Recent discoveries challenge this long-held belief, with the pig brain regaining metabolic and spontaneous neuronal activity several hours after death, although synchronized global activity remains absent [4], [5]. However, efforts to preserve the brain have not exceeded six hours after circulation was restored, leaving the question of whether a more complete recovery might be possible over a longer timeframe.

Complimenting these observations, we recently restored postsynaptic ON-bipolar cell light responses in postmortem human retinas recovered within 20 minutes of circulation loss but not in eyes enucleated 1- 5 hours postmortem [6]. While inner retinal neurons can partially regain spontaneous activity and light responsiveness following brief hypoxia or ischemia [7], [8], [9], [10], synchronized network activity of CNS tissues, including the retina, has not been achieved beyond 30 minutes after circulatory death. Interestingly, partial recovery of inner retinal light responses several weeks after retinal ischemia in a rhesus monkey model of central retinal artery occlusion (CRAO) [8] suggests that retinal ischemic damage may be reversible with a sufficient recovery time.

Current treatments for CRAO center around rapidly restoring blood flow through thrombolysis, ocular massage, paracentesis, intraocular pressure reduction, or hyperbaric oxygen therapy [11], [12], [13]. However, the narrow therapeutic time window and limited efficacy of these treatments result in legal blindness (20/200 visual acuity or worse) in the affected eye of 81% of CRAO patients [14]. Extending the viability of neurons and understanding mechanisms that reverse ischemic damage could dramatically improve clinical outcomes, not only for patients with CRAO but also for other conditions involving acute circulatory loss of the CNS, such as in ischemic stroke.

In this study, we report the recovery of retinal function following prolonged ischemia. By maintaining retinal light responses in the laboratory, developing *ex vivo* infrared OCT and fundus imaging to precisely identify and sample retinal structures in human donor eyes, and a recently developed small-volume closed perfusion system [15], [16], we advance the use of the postmortem human retina as a powerful research tool. Additionally, we present a functional *ex vivo* assay to model diseases associated with acute ischemia-reperfusion injury, such as CRAO, and to evaluate drugs targeting ischemia-reperfusion injury.

## RESULTS

### Restoring and preserving light signaling in human donor retinas

We previously revived photoreceptor and ON-bipolar cell light responses when eyes were recovered within 20 minutes of circulatory arrest (cross-clamp). In contrast, we recorded only photoreceptor light responses if enucleation was delayed by more than one hour [6]. To explore the necessary time window for restoring *in vivo*-like light responses with *ex vivo* ERG, we partnered with the Utah Lions Eye Bank and the Organ Procurement Organization DonorConnect in Salt Lake City, Utah to obtain whole human globes or posterior poles (after collection of the corneas for transplantation) from donations after brain and cardiac death (DBD and DCD, respectively).

Peripheral retinas from five DBD or DCD donors, collected 50-60 minutes after circulatory arrest, showed only small photoreceptor light responses during the first 1-2 hours after arriving in the laboratory (**Fig 1A**). Since ischemic damage can be reversed in models of CRAO [8], we tested whether overnight incubation could improve light responses. Indeed, by incubating eyecups (with the cornea, lens, and some of the vitreous removed) or pieces of the posterior eye in oxygenated bicarbonate-buffered Ames’ media at room temperature overnight (**Fig 1B**), we revived postsynaptic ON-bipolar cell light responses (**Fig 1C, E, F**) and recorded large photoreceptor light responses (**Fig 1D**) in eyes collected within 50-60 minutes. Notably, this protocol also restored ON-bipolar cell responses in 2 out of 5 pairs of research donor eyes enucleated 3-4 hours postmortem (**Supplementary** Fig 1).

**Figure 1.**
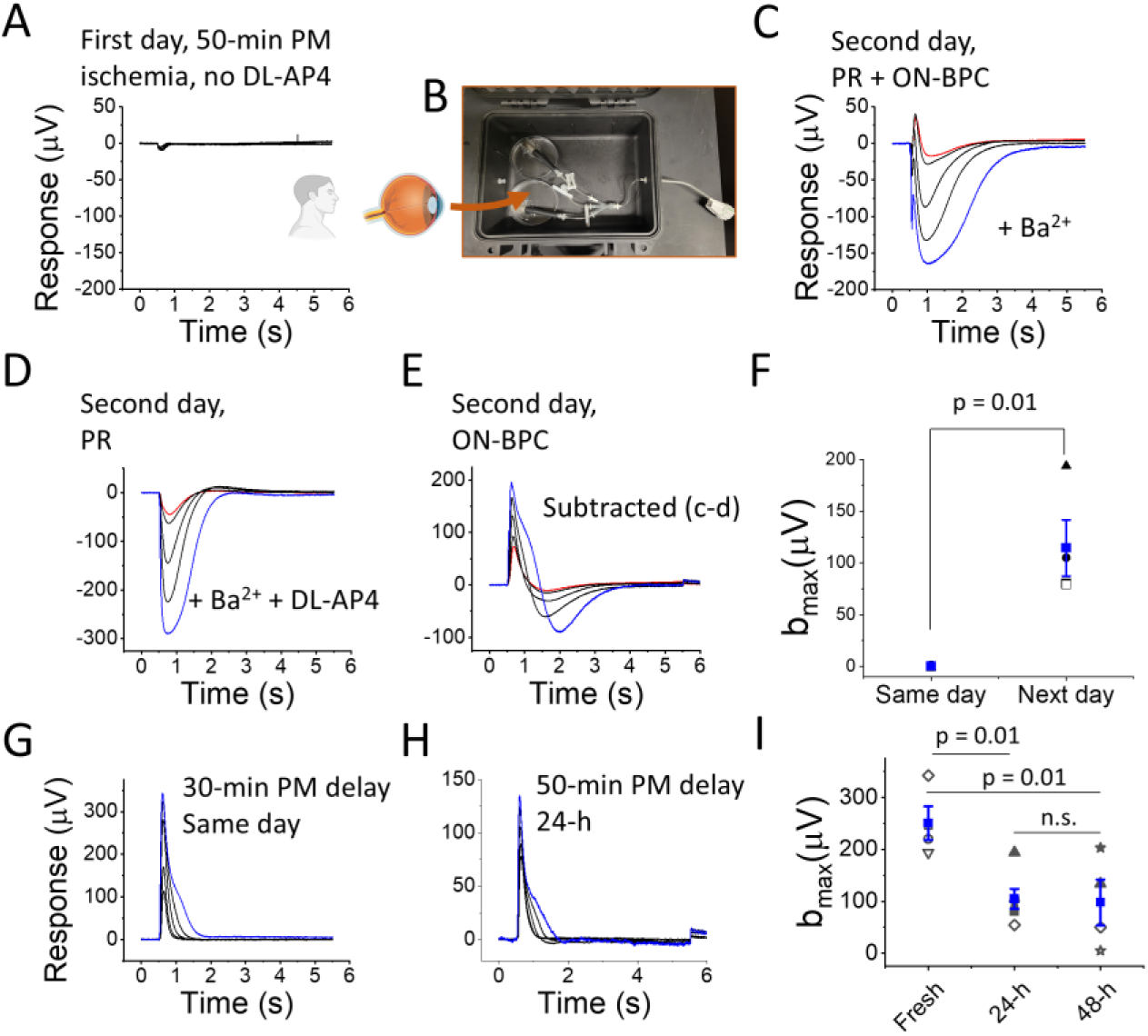
Restoring and preserving ERG b-waves in organ donor eyes after overnight eyecup incubation. (A) *Ex vivo* ERG light responses from the peripheral retina of an eye enucleated 50 min postmortem, recorded on the same day. In the absence of DL-AP4 only small a-waves were observed. (B) Overnight incubation of the eyecup in oxygenated media in darkness. (C) Light responses of a peripheral retina sample (after overnight eyecup incubation) in the presence of Ba^2+^ alone and (D) combined with DL-AP4. (E) Subtracting responses in (D) from those in (C) isolates a strong b-wave. (F) Maximum b-wave amplitudes (i.e., blue trace in (E) from three donors with 50-65-min enucleation delay, recorded ∼3 h postmortem (Same day) or after overnight incubation (Next day) (Blue: mean ± SE). (G) *Ex vivo* ERG b-waves determined by subtraction, as shown in c-e, from human DBD eye enucleated within 30 min after death and recorded on the same day and (H) from human DCD eye enucleated within 50 min after death and recorded following 48 h incubation. (I) Maximal b-wave amplitudes (mean ± SE) from peripheral samples recorded soon after enucleation (fresh, 3 donors with <30 mins enucleation delay), 24 (5 donors with <60- mins enucleation delay) and 48 h (4 donors with <60-mins enucleation delay) after enucleation. Light flash strengths ranged from 19 to 1,014 photons (530 nm) μm^-2^.

It is crucial that the retina remains attached to the retinal pigment epithelium (RPE) during incubation. Isolated mouse retinas lost almost all of their ON-bipolar cell and much of their photoreceptor light responses during overnight incubation, even when kept in oxygenated media (**Supplementary** Fig 2). These findings show that postmortem ischemic injury can be reversed after prolonged incubation of retinas attached to the RPE, provided the eyes are enucleated and placed in oxygenated media within approximately one hour, or in some cases, several hours, from the loss of circulation.

Although human retinal organoids and, on occasion, neural retinas have been cultured long-term [17], [18], [19], [20], [21], [22], [23], preservation of the whole human posterior eye *ex vivo* remains largely unexplored. To determine the duration for which retinal light responses can be maintained, we incubated posterior eyecups in oxygenated bicarbonate-buffered Ames’ media at room temperature, as described above (**Fig 1B**). We punched 5 mm retina samples near the macula at 12, 24, and 48 hours postmortem and determined the subtracted *ex vivo* ERG b-wave amplitude (ON-bipolar cell response) as a sensitive biomarker for viability, as described in **Fig 1C-E**. With good retinal attachment and aseptic incubation media, we preserved ON-bipolar cell light responses for at least 48 hours (**Fig 1G-H**), although the maximum response amplitudes declined over time (**Fig 1I**).

### Visualizing and Collecting Human Retina Samples for *Ex Vivo* ERG Recordings

The macula and fovea, located in the central retina and unique to humans and non-human primates among mammals, mediate high-acuity daytime color vision [24]. Current animal models fail to faithfully recapitulate the physiology and pathophysiology of the central retina, making it challenging to study diseases that affect high-acuity vision, e.g., age-related macular degeneration [25]. Additionally, current technologies do not permit studying macular physiology in postmortem eyes from healthy donors and patients with retinal diseases. To fill this gap, we adapted an existing imaging platform to enable precise sample collection from human donor eyes while preserving retinal light sensitivity.

We converted a Heidelberg Spectralis Optical Coherence Tomography (OCT) and fundus imaging platform for *ex vivo* analysis of eyecups under infrared light (**Fig 2A, B**). Typically used for high-resolution cross-sectional images of the retina in clinical settings, we adapted the OCT to image postmortem eyecups filled with Ames’ media after the cornea and lens were removed to provide a clear optical path. A custom scanner lens adapter was created with Fusion 360, with which we securely fitted an additional lens to correct for altered refraction after removing the lens and cornea (shown in blue in **Fig 2A and C**; non-contact animal imaging lens, +25 Diopter, from Heidelberg Engineering Inc.). This setup allows us to image the retina and accurately locate the fovea in human donor eyes (**Fig 2B**).

**Figure 2.**
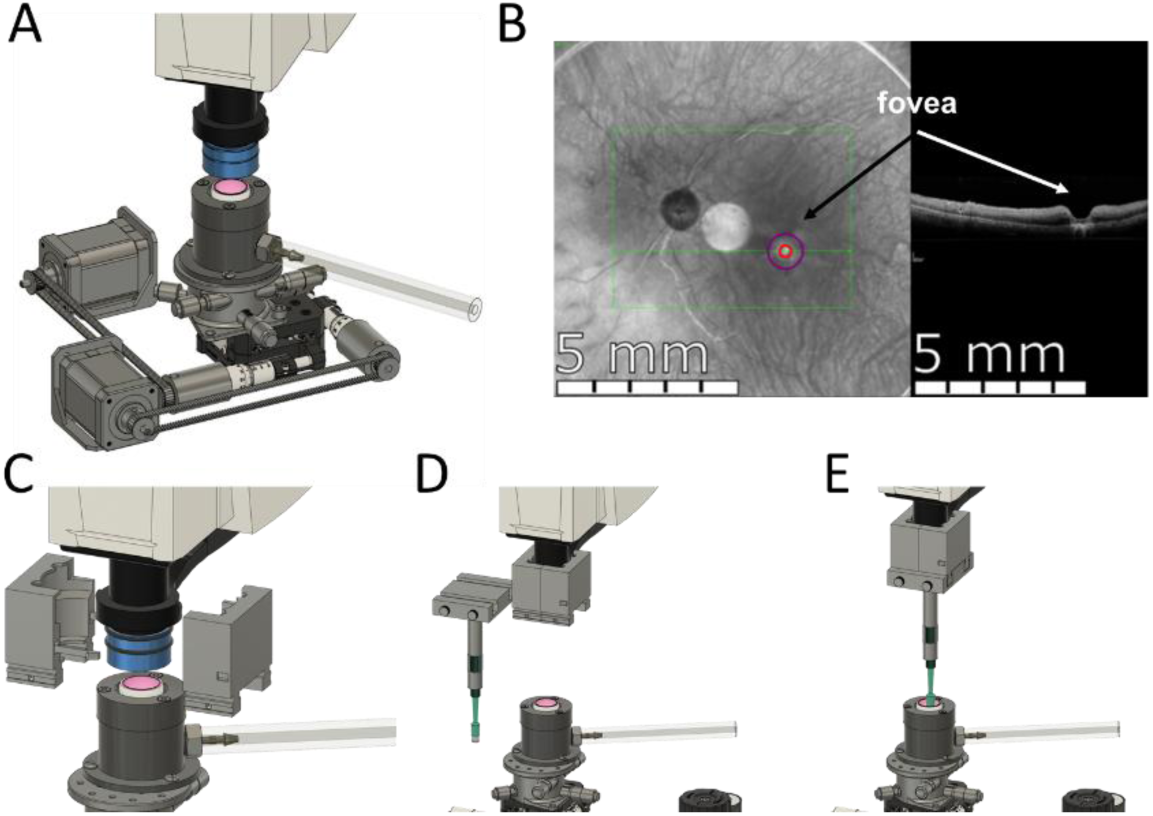
Foveal analysis and procurement automation. (A) Fundus and OCT imaging of *ex vivo* human eyecup. (B) Deep learning models scan the retina, locate the fovea and move the retina into position. (C-D) Biopsy punch attachment affixed to OCT lens. (E) Sampling of the fovea.

A punch mechanism attached to a sliding lock to collect retinal samples (**Fig. 2C-E**) (https://github.com/jordanallen291/Automatic-Foveal-Detection) achieved a puncture localization accuracy of 0.100 ± 0.066 mm across 24 trials (**Supplementary** Fig 3A). Although effective with a 100 % success rate (9/9) under illuminated conditions (**Supplementary** Fig 4), this method was prone to human error under dim red light in our electrophysiology suite, prompting us to develop two deep-learning models that automatically locate the fovea. The first model used a binary classification scheme to determine whether OCT images contain a fovea with 100% accuracy in the training and validation phases (**Supplementary** Fig 3B). The second model, based on the Ultralytics YOLO v8 architecture, precisely located the foveal coordinates with 98% precision and a 90% recall across images. This model effectively identified instances of the target class, ’fovea’, in 30 validation images (**Supplementary** Fig 3C). All code and AI models are available on GitHub (https://github.com/jordanallen291/Automatic-Foveal-Detection).

Using the described fundus- and OCT-imaging, we collected the foveal pits and surrounding maculae (**Fig 3A**). Light responses from a DBD donor (who had been on life support for four days) eyes, which were enucleated 30 minutes after cross-clamp, showed strong photoreceptor and downstream retinal neuronal light responses in the central and peripheral retina regions (**Fig 3D, C, and G, F**, respectively). This is consistent with our previous work which revived robust outer and inner retina light responses when postmortem enucleation delays were between 0 to 20 minutes [6].

**Figure 3.**
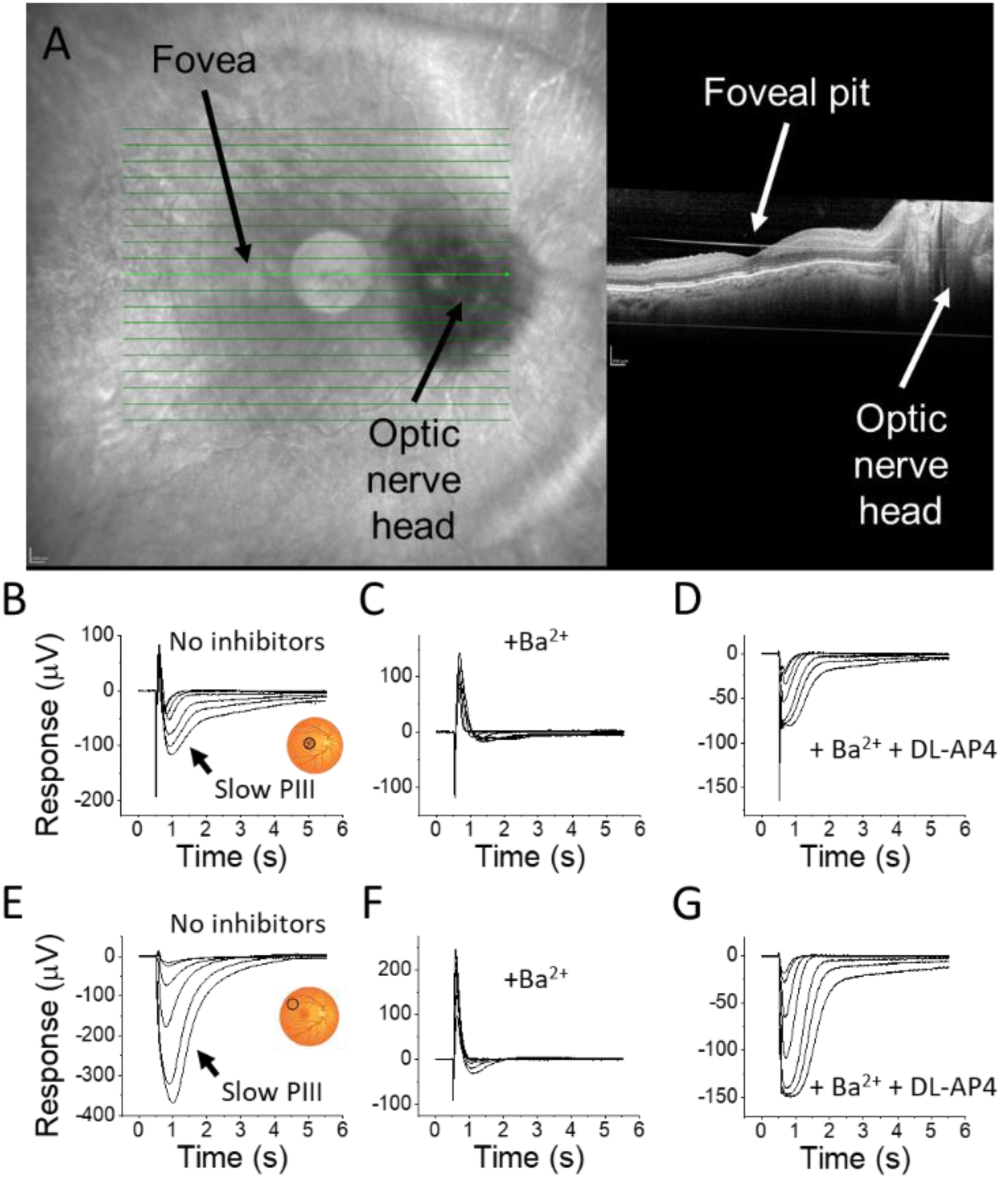
Imaging and recording of light responses from the central and peripheral human retina. (A) Infrared fundus and OCT image of the postmortem human retina from an eye enucleated 30 min postmortem. Retinal light responses in the central (B-D) and peripheral (E-G) retina from a DBD eye. Macular and peripheral light responses from the same eye shown in (A), (B, E) without inhibitors, (C, F) with 100 µM BaCl_2_ to block glial responses and (D, G) with BaCl_2_ and 40 µM DL-AP4 to remove glial and inner retina components. Light flash strengths at 530 nm wavelength ranged from 19 to 3,525 photons μm^-2^.

In addition, we are the first to identify a Ba^2+^-sensitive slow negative wave, known as the slow PIII (glial) component, in the macular region of the human retina (**Fig 3B**). The slow PIII wave has been extensively described in cold-blooded species and rodents and likely arises from K^+^ buffering by Müller glial cells [26], [27], [28], [29]. In the human peripheral retina, the slow PIII component was noticeably larger and similar to that in mice, compared to the smaller Ba^2+^-sensitive component in the macula (**Fig 3B, E**).

### Drug Testing in the Human Retina and Development of an Ischemia-Reperfusion Injury Model

To create a drug testing platform that minimizes drug usage [15], [16], we built a closed perfusion system that recirculates small perfusate volumes (150-180 mL) as recently described [15]. After overnight incubation of three human donor eyecups with 1–4 hour enucleation delays, peripheral retina samples maintained stable *ex vivo* rod photoreceptor light responses for at least 12 hours after initiating perfusion in the *ex vivo* ERG specimen holder. During this time, the dim flash response time-to-peak and amplitude, the normalized amplitude to the maximal response, the maximum response, the normalized maximum response to the initial response, and the flash strength producing half-maximal responses remained stable (**Fig 4**). Consistent results from all three donor retinas confirmed that our setup preserves retinal function during extended recordings with a minimal perfusate volume.

**Figure 4.**
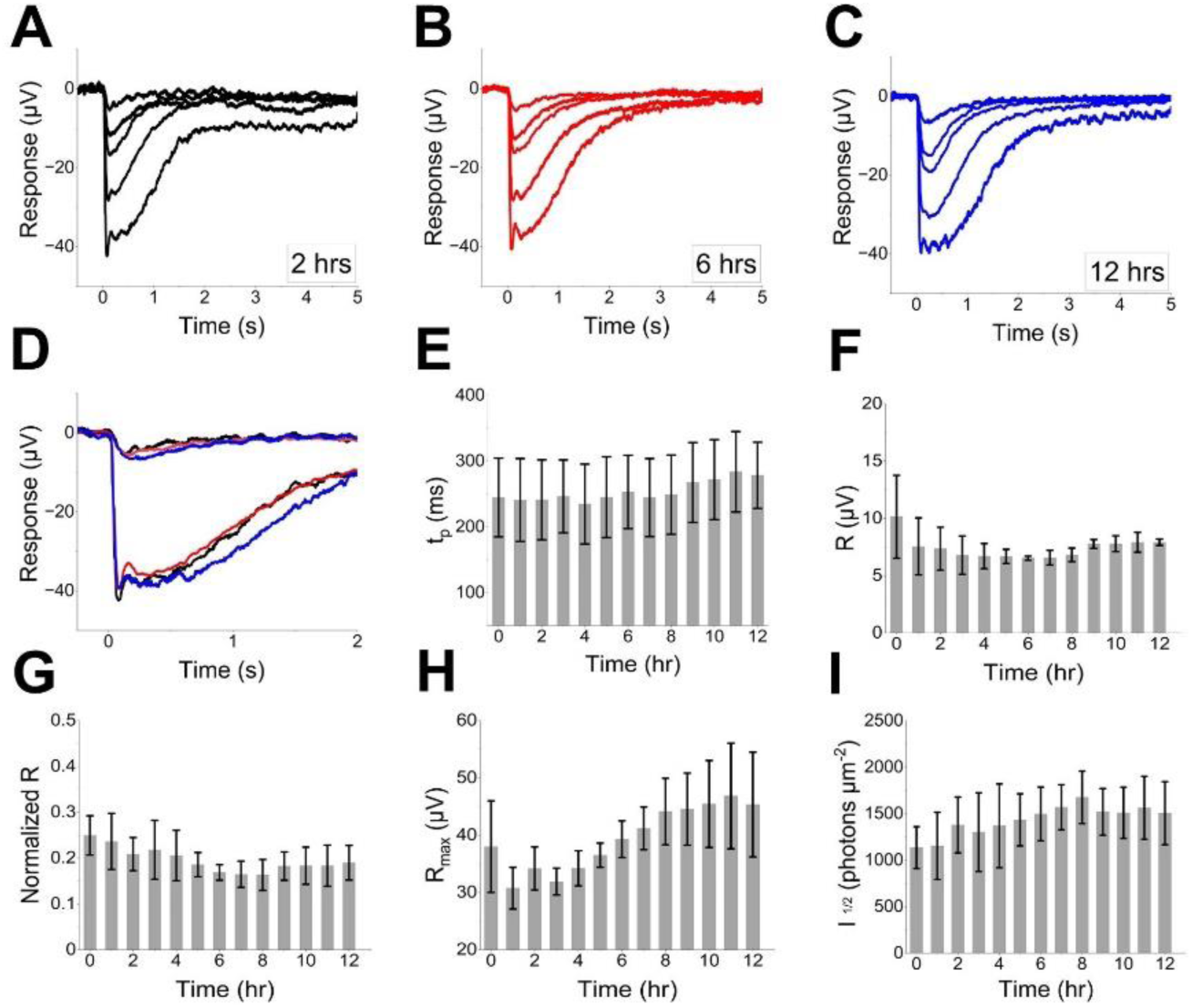
Preservation of ERG a-waves in the human retina during small-volume closed perfusion. (A-C) Example traces from a 38-year-old anoxic brain injury case with a one-hour post-mortem enucleation delay. (D) Preservation of dim and bright photoreceptors light responses at 2 (black), 6 (red) and 12 hours (blue) after perfusion was started. (E) time to peak, (F) dim stimulus amplitude (R), (G) normalized amplitude to the maximal response, (H) maximum response and (I) light sensitivity (I_1/2_). Data in E-I is from three donors (UFT8, D54_2 and D55, see table 1), mean±SEM.

### Acute oxygen deprivation in the mouse and human retina as a model for ischemia/reperfusion injury

Several *in vivo* animal models study ischemia-reperfusion injury in the eye and brain caused by, e.g., cardiac arrest, stroke, and central retinal artery and vein occlusion [3], [8], [30], [31], [32]. However, no *ex vivo* models exist to assess retinal function following ischemia-reperfusion injury in postmortem human retinas. To address this gap and expand the use of our *ex vivo* testing platform, we developed an acute ischemia-reperfusion injury model that replicates neuronal damage caused by hypoxia and reoxygenation. This model is based on hypoxia/reoxygenation, since our previous work demonstrated that postmortem loss of retinal light responses is likely caused by hypoxia, not tissue acidification due to ischemia [6].

To determine the relative susceptibility of different retinal neuronal cell types to hypoxia we superfused isolated mouse retinas with hypoxic Ames’ media (∼2.5% O_2_) while recording *ex vivo* ERG. We observed that ON-bipolar cells lost their light responses within several minutes while photoreceptors retained >50% of theirs for at least 30 minutes (**Supplementary** Fig 5). The greater vulnerability of ON-bipolar cells to hypoxia compared to photoreceptors is also supported in patients with CRAO [33]. However, CRAO predominantly affects the inner retinal perfusion while choroidal perfusion is maintained, offering an alternative explanation for the greater vulnerability of the inner retinal neurons.

We determined the time course of ischemia/reperfusion injury by measuring light responses in mouse eyecups exposed to low oxygen concentrations: Light responses were substantially reduced immediately after one and three hours of hypoxia, with longer hypoxia resulting in greater loss of ON-bipolar cell light responses (**Fig 5A-C**). Overnight incubation in oxygenated Ames’ media fully restored photoreceptor and ON-bipolar cell function after one hour of hypoxia, but only partially restored function after three hours of hypoxia (**Fig 5A-C**). When we superfused the retina in the specimen holder with hypoxic Ames’ media for one hour (by bubbling the perfusion media with 95% N_2_ and 5% CO_2_) and simultaneously recorded light responses, ON-bipolar cell light responses were eliminated (**Fig 5E**). However, in two out of three samples, these responses substantially recovered within 15 minutes of reoxygenation (**Fig 5D, E**).

**Figure 5.**
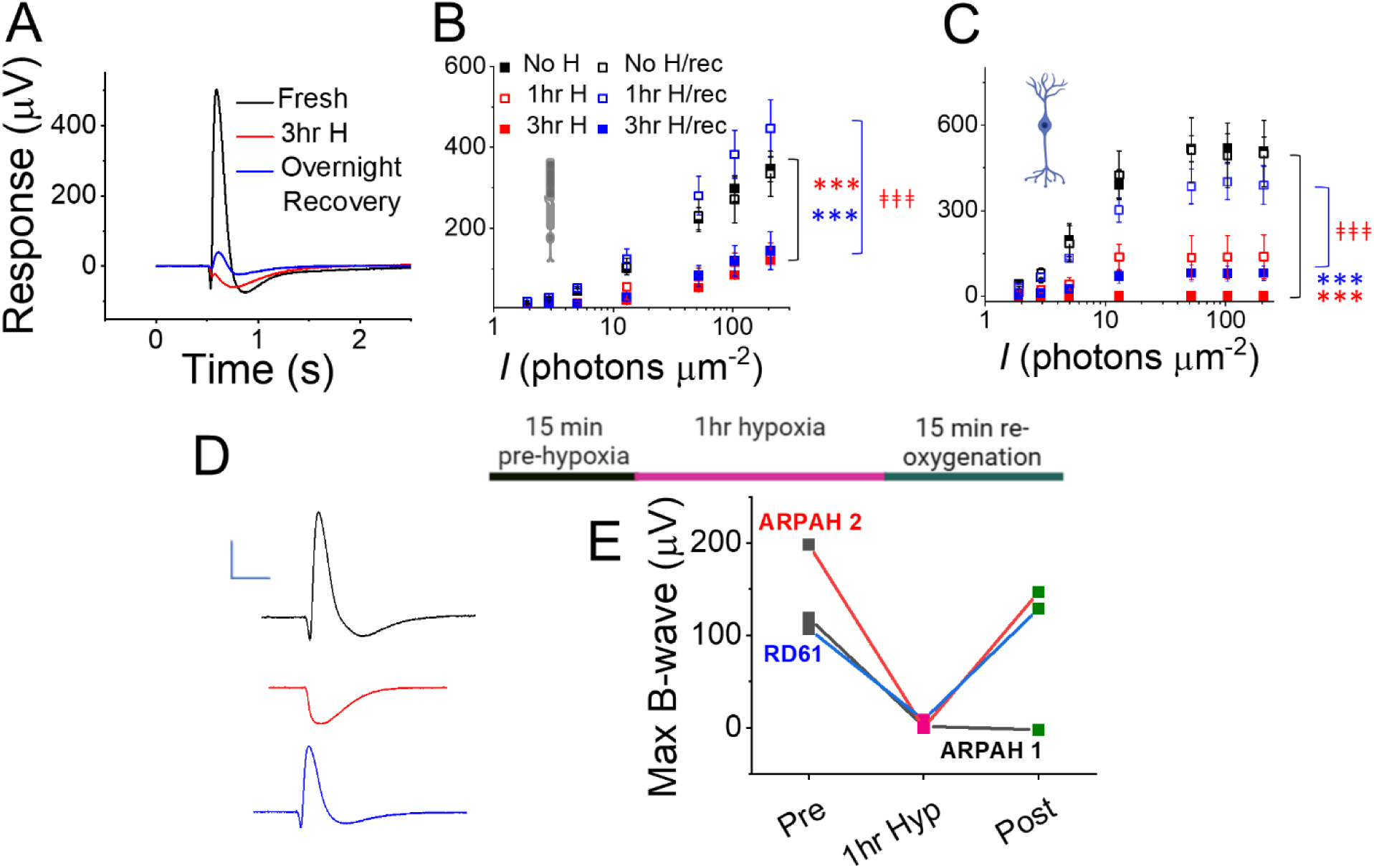
Ischemia-reperfusion injury model in the mouse and human retina. (A) Example traces showing rod photoreceptor light responses in *Gnat2* knockout mouse retinas immediately after death (black), after three hours of hypoxia (red), and after three hours of hypoxia followed by overnight recovery in oxygenated media (blue). (B) Averaged *Gnat2* knockout mouse rod photoreceptor and (C) ON-bipolar cell responses after one or three hours of hypoxia with and without overnight recovery in oxygenated media compared to retinas kept in oxygenated media without hypoxia (n = 2-6). *** P<0.001 (compared to oxygenated control), ‡‡‡ P<0.001 (compared to hypoxia). (D) Example traces showing light responses in human donor retina in oxygenated media (black), after perfusion with hypoxic media in the *ex vivo* specimen holder for one hour (red), and after recovery in oxygenated media for 15 minutes (blue). Scale bars: 100 µV (y) and 1 s (x). (E) Averaged ON-bipolar cell light responses from three human donor retinas under the same conditions as in (D).

### Mechanisms of ischemic damage and repair

Ischemia-reperfusion injury causes multi-factorial damage, including an oxidative stress response from NADPH oxidase and the mitochondrial electrode transport chain. Reduced ATP production during hypoxia impairs intracellular Ca^2+^ extrusion and increased intracellular Ca^2+^ activates NADPH oxidase [34]. The primary reactive oxygen species (ROS) produced by NADPH oxidase, superoxide, and H_2_O_2_, open mitochondrial permeability transition pores, uncouple the mitochondrial electron transport chain, and further increase superoxide production [35]. In addition, succinate accumulates, e.g. in the ischemic heart [36], which after reperfusion generates ROS by reversing complex I of the mitochondrial electrode transport chain.

To counteract this damage we tested pharmacological agents: Dimethyl malonate (DMM), a succinate dehydrogenase blocker that prevents succinate accumulation during hypoxia and reduces ROS production in the mitochondrial electron transport chain during reoxygenation [37], and apocynin which blocks the assembly of the NADPH oxidase [38]. Neither drug prevented the immediate loss of retinal light responses after one hour of hypoxia (**Fig 6A, B**). However, apocynin improved overnight recovery of photoreceptor, but not ON-bipolar cell, function after three hours of hypoxia (**Fig 6C, D**). DMM showed a non-significant trend toward improved photoreceptor function (**Fig 6C, D**). This implies that excessive ROS production during reoxygenation impairs photoreceptor, but not ON-bipolar cell responses.

**Figure 6.**
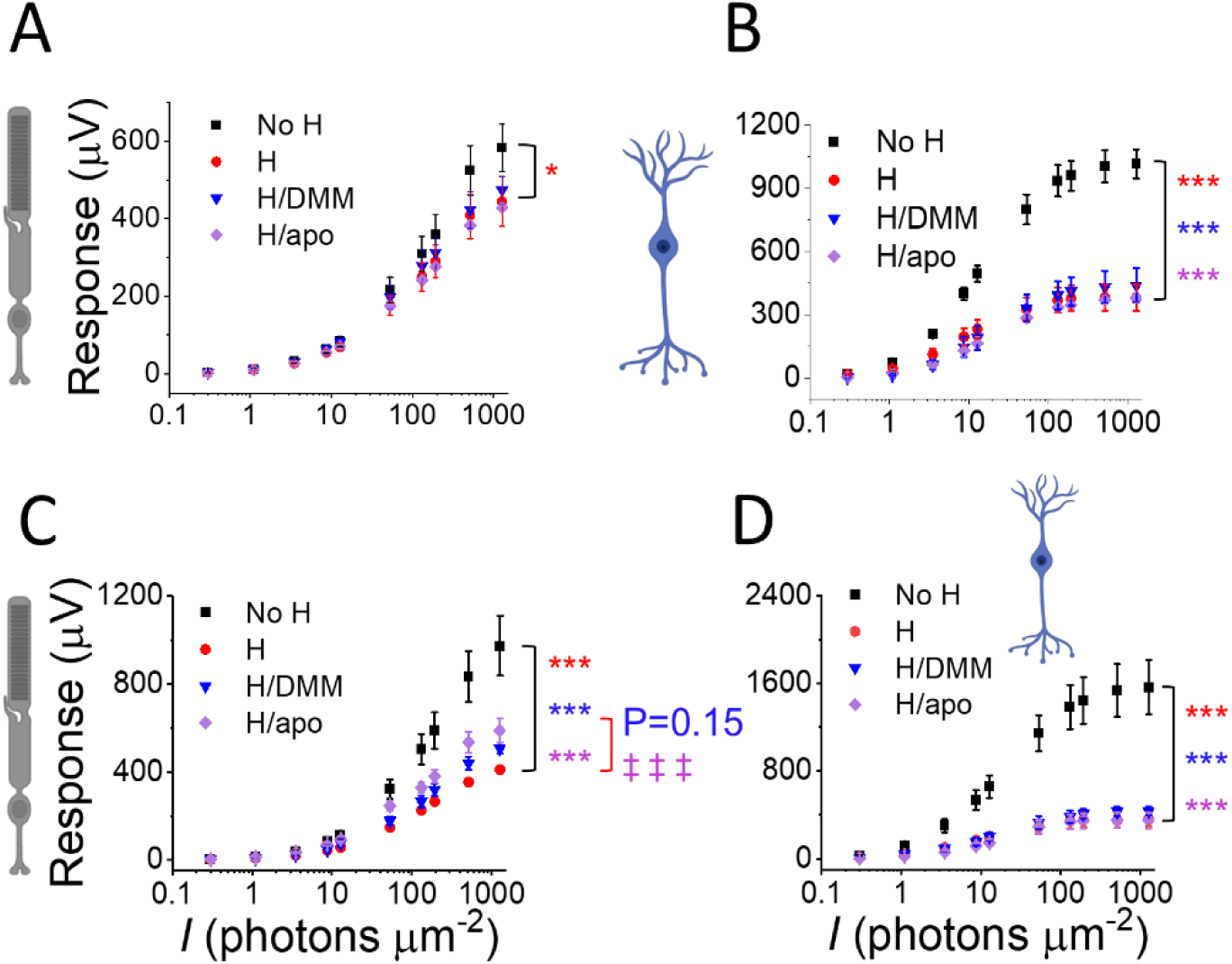
Preventing retinal injury during hypoxia and reoxygenation. Photoreceptor (A, C) and ON-bipolar cell light responses (B, D) in oxygenated media or (A, B) immediately after one hour of hypoxia (H) or (C, D) after three hours of hypoxia (H) with overnight incubation in oxygenated Ames’ media in the absence or presence of dimethyl malonate (DMM) or apocynin (apo, n=4-6). * P<0.05, *** P<0.001 (compared to oxygenated control), ‡‡‡ P<0.001 (compared to hypoxia).

By carefully timing and combining these inhibitors we revealed more of the underlying repair mechanisms: DMM most effectively protected photoreceptors when administered during hypoxia or just before reoxygenation (**Supplementary** Fig 6), suggesting that succinate accumulation during hypoxia leads to a detrimental burst of ROS by reversing succinate dehydrogenase immediately upon reoxygenation. Applying DMM only during reoxygenation did not protect photoreceptor function (**Supplementary** Fig 6). Interestingly, combining DMM and apocynin was less protective for photoreceptors than apocynin alone, while the general ROS scavenger α-lipoic acid impaired photoreceptor recovery (**Supplementary** Fig 6). None of these pharmacological agents alone or in combination was protective for ON-bipolar cells (**Supplementary** Fig 6). These findings suggest that excessive ROS production by the electron transport chain and the NADPH oxidase may not be the main mechanisms of damage to ON-bipolar cells, but damages photoreceptors. Limited ROS production may be necessary to prevent or repair ischemia-reperfusion injury in both cell types.

Finally, we tested two NMDA receptor blockers, MK-801 and memantine, to prevent excitotoxicity in inner retinal neurons caused by excessive glutamate release from damaged photoreceptors and ON-bipolar cells under hypoxic conditions and during reoxygenation. Neither drug was protective, and MK-801 was detrimental to light responses after three hours of hypoxia (**Fig 7**).

**Figure 7.**
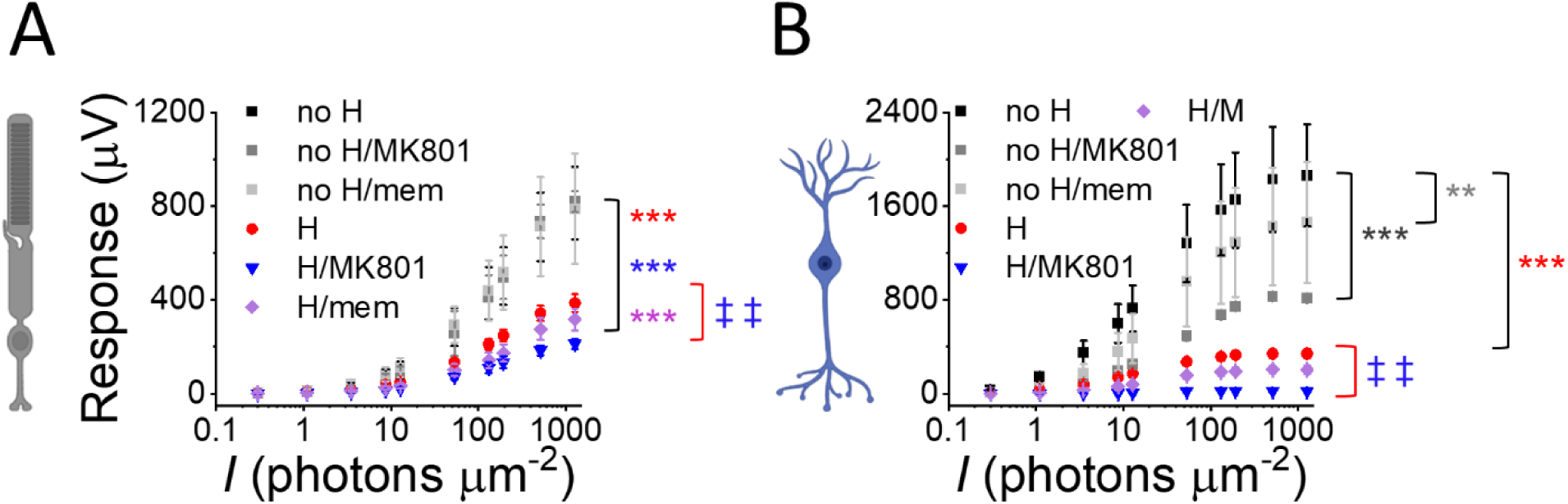
NMDA receptor blockers do not prevent retinal injury during hypoxia and reoxygenation. Photoreceptor (A) and ON-bipolar cell light responses (B) in oxygenated media (O) in the absence or presence of MK-801 or memantine (mem) or after three hours of hypoxia (H) with overnight incubation in oxygenated Ames’ media in the absence or presence of MK-801 or memantine (n=2-6). ** P<0.01, *** P<0.001 (compared to oxygenated control), ‡‡ P<0.01 (compared to hypoxia).

## DISCUSSION

Despite extensive animal research, more than 90% of clinical trials for investigational new drugs (INDs) fail to progress to market approval [39], [40], often due to inadequate safety and efficacy data from animal models [25], [41], [42], [43], [44]. Small animal models and in vitro assays frequently struggle to replicate the complex pathologies of retinal diseases, e.g., age-related macular degeneration (AMD) and glaucoma, which are influenced by advanced age, multiple genetic risk factors of low penetrance, and environmental and lifestyle factors [25]. Non-human primates share key retinal features with humans, but current disease models in these species are limited to certain forms of inherited retinal degeneration [45], and their use is constrained by ethical concerns and high costs [25].

Research in human postmortem eyes, especially those from donors with retinal diseases, holds immense potential to overcome these limitations. Such research could provide valuable insights into disease mechanisms and facilitate the development of new treatments, offering hope to millions of patients facing vision loss and diminished quality of life. Several challenges have previously restricted the use of human postmortem eyes for large-scale physiological experiments, including the need for ultra-fresh human tissue, the difficulty of preserving light responses in the laboratory, and challenges in sampling precise retina locations.

We address these barriers by establishing a robust infrastructure in collaboration with the Utah Lions Eye Bank and DonorConnect, enabling regular, weekly recovery and transportation of light-responsive human eyes from DBD and DCD donors (**Figs 1, 3**). By scaling access to functionally viable human tissue, we lay the foundation for detailed studies of human retinal physiology, neural network activity, and pharmacological testing.

Our previously published data clearly showed that mouse and human photoreceptors maintain their light responses more effectively during postmortem ischemia and better recover their function immediately after hypoxic injury compared to ON-bipolar cells [6]. Here, we report that ON-bipolar cell light responses, particularly in the peripheral retina, can be restored by overnight incubation of eyecups in oxygenated Ames’ media, even after 60 minutes of postmortem ischemia or hypoxia (**Fig 1, 6**). Peripheral rod ON-bipolar cell responses were sometimes even restored in research donor eyes enucleated 3-4 hours postmortem, although their responses remain smaller than those recorded from freshly recovered organ donor eyes, and photoreceptor light response amplitudes consistently improved (**Supplementary** Fig 4). Importantly, light responses of central retinal ON-bipolar cells remained elusive, suggesting that the macula and/or cone pathways may be more susceptible to ischemic injury. Since central vision loss has the greatest impact on patients, our future studies will focus on optimizing methods to preserve or restore ON-bipolar cell function in the central retina.

Our findings challenge the widely held notion that ischemic or hypoxic injury irreversibly damages neurons, including those in the retina. Instead, they suggest that significant neuronal recovery is possible with timely restoration of perfusion (**Fig. 1, 5**). This aligns with a recent study reporting cellular recovery in pig brains following one hour of warm ischemia [5]. These data imply that, instead of collectively undergoing hypoxia-induced apoptosis, at least a sizeable proportion of retinal neurons survive. Temporary loss of ON-bipolar cell light responses must, therefore, be attributed to loss of synaptic input from photoreceptors and/or ischemia-induced membrane potential dysregulation, preventing light signal transmission in ON-bipolar cells.

However, irrespective of the mechanism(s) underlying the transient loss of ON-bipolar cell function, we importantly not only extend access to functionally viable human eyes but also question whether ischemic or hypoxic retinal or brain injury is truly irreversible and can be repaired as long as cells are reperfused within a critical time frame. Even though we consistently restored and preserved cone photoreceptor light responses in the maculae from DCD and research donors, and rod-mediated photoreceptor and ON-bipolar cell light responses in the peripheral retina, signal transmission to cone ON-bipolar cells remained elusive, indicating that the central retina and/or cone pathway may be more susceptible to ischemic injury. Our future research should, therefore, identify improved methods that consistently retain or restore ON-bipolar cell light responses in the central retina.

*Ex vivo* ERG in human donor retinas overcomes many disadvantages of alternative techniques, particularly when studying age-related physiological changes and multifactorial diseases. Unlike mice with their shorter lifespans and retinal organoids, which largely replicate retinal development [17], [25], our platform provides access to mature human tissue. Non-human primates, while anatomically similar to the human retina, including a macula, lack relevant disease models and face limitations due to high costs and ethical concerns. Our approach provides a practical, human system to study central vision loss, which cannot be effectively studied in non-primate mammals.

Our study provides protocols and technologies for long-term functional preservation of eyecups, precise sample collection, mechanistic studies, and drug testing (**Fig 1-5**). Using these methods we investigated pathways involved in ischemia-reperfusion injury in the retina with a focus on oxidative stress and glutamate excitotoxicity (**Fig 6-8**). These experiments highlighted that inhibiting specific ROS-producing pathways, such as succinate dehydrogenase (Complex II) and NADPH oxidase, can protect photoreceptor function during and after hypoxia. However, combining ROS inhibitors yielded less protection than expected, and broad-spectrum scavenging of ROS with α-lipoic acid was detrimental, suggesting that limited ROS may play protective roles. Furthermore, while photoreceptor function was improved by limiting ROS, that of ON-bipolar cells was not (**Fig. 6**), indicating that different protective strategies may be required for different retinal cell types.

Our findings also have implications for developing novel treatments for acute retinal ischemia, e.g., in CRAO. Current treatments, including hyperbaric oxygen therapies and intraocular pressure reduction, are often insufficient, and patient care lacks effective treatment options. Our *ex vivo* data suggests that supplying the retina with oxygen within a critical time frame of less than three hours can dramatically benefit light responses in retinal neurons. This raises the possibility that vitrectomy and perfusion of the vitreous cavity with oxygenated media, possibly in combination with ROS-targeting or neuroprotective drugs, could be developed as a therapeutic intervention for CRAO to mitigate ischemia-reperfusion injury until retinal perfusion is restored.

Implications of our study may even extend beyond ophthalmology, potentially offering insights into ischemia-reperfusion injury in other CNS tissues. The retinal ischemia-reperfusion model may inform treatment strategies for acute damage following ischemic stroke or brain injury after cardiac arrest. Additionally, our platform will bridge the gap between in vitro and clinical testing of INDs, providing human-relevant information that may correlate with clinical outcomes.

Finally, our work will support efforts to replace retinas or even whole eyes, similar to the pioneering whole-eye transplantation recently performed at NYU Langone Health [46], [47] and the audacious program announced by the Advanced Research Projects Agency for Health (ARPA-H) to restore vision in bilaterally blind patients through whole-eye transplantation (THEA) [48]. Our methods for recovering, restoring, and preserving light responses in human donor eyes are directly applicable to these efforts by supporting light responses in the central and peripheral retina from DBD and DCD organ donors.

## Acknowledgements

We thank Austin Adderley, Lana Gale, Wade McEntire, and the recovery technicians at the Utah Lion’s Eye Bank, Brent Davis and the Donor Connect Surgical Recovery Coordinators team, as well as the University of Utah and Intermountain transplant surgeons for their support in procuring human donor eyes. This research was, in part, funded by the Advanced Research Projects Agency for Health (ARPA-H). The views and conclusions contained in this document are those of the authors and should not be interpreted as representing the official policies, either expressed or implied, of the United States Government. This work was also supported by NEI R01 EY031706 to F. V., as well as a DRC Research grant to S. B., National Institutes of Health Core Grant (EY014800), and an Unrestricted Grant from Research to Prevent Blindness, New York, NY, to the Department of Ophthalmology & Visual Sciences, University of Utah. Also, Dr. Frans Vinberg is a recipient of a Research to Prevent Blindness Dr. H. James and Carole Free Career Development Award. This work was also supported in part by the John A. Moran Eye Center (JMEC CORE) under Grant P30 EY014800 and Vision Research Training Grant T32 EY024234.

## Methods

### Donor Demographics, Eye Procurement and Transportation

Human eye tissue was procured by the Utah Lion’s Eye Bank (Food and Drug Administration accreditation number 3000215349) in conjunction with whole organ donation by Donor Connect (Food and Drug Administration accreditation number 3000215350, Centers for Medicare & Medicaid Services accreditation number 46P001). The Utah Lion’s Eye Bank obtained consent for research use for all donor eyes according to the Uniform Anatomical Gift Act (UAGA). The research team did not identify or select potential organ donors and had no access to non-anonymized donor information, except for information on ocular diseases, such as age-related macular degeneration and diabetic retinopathy. Our research project was, therefore, classified as exempt by the University of Utah (IRB no. 00106658).

Whole eyes or posterior poles were enucleated within 30-60 minutes after cross-clamp and transported to the laboratory in HEPES-buffered Ames’ media that was directly bubbled with 100% O_2_ at room temperature in the custom transportation case we previously described [6].

Eyes from research donors were enucleated up to 5 hours postmortem by a trained Eye Bank technician and immediately placed into HEPES-buffered Ames’ media at room temperature. Eyes from organ donors were obtained up to 1 hour after cross-clamp at the time of organ removal, either with or without corneas, if these were transplanted, and placed into HEPES-buffered Ames’ media and oxygenated with 100 % medical oxygen in our custom transportation case. From the time of retrieval, eyes were protected from light, stored at room temperature, and immediately transferred to the laboratory. Table 1 lists basic non-identifying information: age, sex, cause of death, cross-clamp/death-to-preservation delay, and potentially relevant medical conditions of all organ and research donors used in this study.

### Sample Preparation

Upon arrival in the laboratory, eyes were handled under dim red-light illumination and stored in bicarbonate-buffered Ames’ media oxygenated with 95 % O_2_ and 5 % CO_2_. By making an incision approximately 3-5 mm behind the limbus, we removed the anterior section of the eye with the lens, if applicable, and removed most of the vitreous to image the retina.

### Hardware

#### OCT Adaptation and Sample Acquisition

We vertically reoriented an optical coherence tomography (OCT) machine (Spectralis, Heidelberg Engineering, Franklin, MA) to ensure that dissected eyecups would remain in place during imaging and sampling. With Fusion 360 (Autodesk, San Rafael, CA), we machined an adapter plate from 6061 aluminum and secured the OCT scanner to additional mounting equipment (ThorLabs, Newton, NJ) uprightly. This and all other custom-made models and mechanical drawings are available on GitHub (https://github.com/jordanallen291/Automatic-Foveal-Detection).

We secured the eyecup in a custom-made vacuum chamber, which we had modeled in Fusion 360 and machined in 6061 aluminum. When we applied pressure of 1.773x10^4^ Pa to porcine eyecups, they remained firmly in position with no discernable signs of external stress. Our custom fixture stage included a LX20/M translational state (X ±25 mm, Y ±25 mm), VAP4/M vertical translation stage (Z ±101.6 mm) and TTR001/M tip, tilt and rotational stage (roll α ±5°, pitch β ±5°, yaw γ ±10°, Thorlabs, Newton, NJ), which provided six degrees of freedom to position the eyecup precisely. After locating the fovea with OCT, we punched a 5 mm retina sample with a disposable biopsy punch (Fisher Scientific, Hampton, NH) on a 3D- printed lens attachment. Elastic O-rings (McMaster-Carr, Elmhurst, IL) reliably and safely secured the sliding biopsy mechanism to the OCT lens.

#### Positional Accuracy Validation

To validate the precision of the punch attachment, we marked twenty-four circles (∼3 mm in diameter) as targets onto an affixed sticky note (3M, St. Paul, MN). A needle puncture outside the targets was marked with tape as the “reference area” before each target was translated to this location and punctured. The distance from the center of each hole to the target was recorded and reported as the mean difference and standard deviation. We used porcine eyecups to test the system’s ability to excise retina samples accurately. To prepare the eyes for testing, we made a small incision near the limbus, submerged the eyes overnight in 4% paraformaldehyde in phosphate-buffered saline, and removed the anterior segment and most of the vitreous.

Using a 2 mm disposable biopsy punch, we made a minor impression in the retina to serve as a reference marker, whose location we marked on the OCT display to compensate for the offset between the OCT’s optical axis and the axis of the biopsy punch. We then aligned the target area with the calibration marker and penetrated the eyecup with the biopsy punch to the sclera. The procedure was deemed successful if the target was accurately incised. This process was repeated three times in three different eyes, resulting in nine data points (N=9).

### Software

We combined several Python scripts to automatically collect retinal images and apply deep learning models that classify images and detect objects. The main program, auto_sample.py, relies on capture_coordinates.py, ImgClassrevised.py, train_obj_det.py, and test_obj_det.py scripts.

#### Screen Coordinate Capture

In the automated workflow of auto_sample.py, the capture_coordinates.py script allows the precise capture of spatial data to interact with the OCT software by interactively collecting screen coordinates and color information through mouse clicks. These coordinates detect when the OCT software is ready for image acquisition and execute a sequence of mouse clicks that will save the acquired images for further analysis. Upon execution, the user is prompted to specify the number of coordinates to be saved. By utilizing both pyautogui and pynput.mouse libraries, the program listens for mouse click events. For each click, it captures and stores the corresponding screen location and the color at that point, leveraging pyautogui.screenshot().getpixel() to obtain the color information. This process documents the selected regions, allowing for automated mouse movement in auto_sample.py.

#### Image Classification

We trained the classification model in auto_sample.py using ImgClassRevised.py to sort OCT images into two categories, either with or without a fovea. The script initially sets up the environment and imports necessary libraries, including TensorFlow, to build and train the neural network and OpenCV for image manipulation. TensorFlow’s image_dataset_from_directory function loads and labels ’fovea’ and ’no fovea’ images from predefined directories based on their folder structure, resizes the images to reduce the computational power needed to train the model, and places the images into batches of 32. The classification model was trained on 592 OCT images from human donor eyes that we had acquired under the same protocol before AI integration, evenly split between ‘fovea’ or ‘no fovea’ categories.

Most of the script preprocesses images by applying random brightness, contrast, saturation, and hue adjustments to optimize the model. The model’s architecture is constructed using TensorFlow’s Keras API, starting with a convolutional base comprising four convolutional blocks, each followed by max pooling to distill features from OCT images efficiently. The initial block introduces 32 3x3 filters with ReLU activation, progressively increasing the filter count to 64 in the second, 128 in the third, and 256 in the fourth to capture a wide array of image features. After flattening the output to a single dimension, the model integrates two dense layers with 512 and 256 neurons, respectively, employing ReLU activation to process the features further. In the final step, a single neuron with a sigmoid activation function designed for binary classification discerns images from those with or without a fovea.

The dataset was divided into training, validation, and test sets to evaluate the model’s performance accurately. Callbacks in the script save the best-performing model with the highest validation accuracy and log training progress, thus facilitating model tuning and evaluation. After training, the script plots loss and accuracy metrics over 30 epochs to visualize the learning process. Finally, users can save the trained model for future use.

#### Object Detection

The train_obj_det.py script delineates the training process of our object detection model and allows auto_sample.py to detect the location of the fovea in classified OCT images with the latest YOLOv8 architecture. The script employs the Ultralytics YOLO library to train a new YOLO model based on a predefined configuration file (yolov8n.yaml), which points to a separate training dataset of 261 annotated foveal images, collected as before with the ImgClassRevised.py script. The train_obj_det.py script ensures the use of GPU if available for faster processing. Its core function, locate_fovea, takes an image path and the YOLO model as inputs and runs inference on the image to predict the presence of a fovea and plots and displays its location. A matplotlib library visualizes the results and uses the PIL library for image processing. After training over 30 epochs, the script validates the model’s performance using a separate set of 30 foveal images.

#### Automatic Sampling

With the data and models from the other scripts, auto_sample.py automatically locates and samples the foveal region of the retina. It initially monitors a predefined screen region to match a target color, indicating that the operator has obtained several b-scans of the retina, and saves the images from the OCT software. The images are each assigned a unique identification number to avoid overwriting existing data, which enables easy access for subsequent processing. The pre-trained script iterates through the saved images, applies necessary preprocessing, and classifies images that contain a fovea. Although the script may find some false-positive images in the peripheral retina, it retains only the longest sequence of positive images, thus indicating the true location of the fovea. The central image of this sequence is kept, as it provides the best representation of the central foveal region.

To obtain the coordinates of the fovea relative to the reference area recorded earlier, the script precisely locates the green b-scan line as the y-coordinate through color space conversion, masking, and edge detection techniques, followed by a Hough Line Transform. Since the scan line on the fundus image is a 1:1 match for the adjacent OCT image, the program can gather the x-coordinate of the fovea based on its coordinates in the OCT image. The script first calibrates the X and Y stepper motors via the Telemetrix library by identifying the difference in the fovea’s location after moving the stage a set distance. Subsequently, it moves the motors to position the fovea directly underneath the biopsy punch for sample acquisition.

### Sample Handling, Preservation, and *Ex Vivo* Electroretinogram

Trained Utah Lion’s Eye Bank technicians recovered human donor eyes, which they immediately submersed in HEPES-buffered Ames’s media (Sigma-Aldrich, St. Louis, MO), protected from light, and transferred to the laboratory [6]. In the laboratory, the tissue was processed under dim red-light illumination, and the donor eyes were immediately transferred into bicarbonate-buffered Ames’ media, which was oxygenated with 95% O_2_ and 5% CO_2_. The anterior segment and most of the vitreous was removed, and the retina was imaged using infrared light and OCT (Spectralis, Heidelberg Engineering, MA). To store eyecups or retinal punches attached to the underlying RPE/choroid until *ex vivo* ERG or for long-term incubation to recover light responses following postmortem delay to enucleation, human eyecups and punches were placed in a deep glass dish filled with bicarbonate-buffered Ames’ media, which was directly bubbled with 95% O_2_ and 5% CO_2_ in darkness.

Mice were sacrificed with CO_2_ asphyxia followed by cervical dislocation. Eyes were immediately enucleated and dissected by piercing the cornea with a 30 G needle, removing the cornea and iris while leaving the lens in place to ensure we did not disrupt the close interaction between the retina and underlying RPE. We stored mouse eyecups in a 100 mL Pyrex bottle filled with Ames’ media, which we either exposed to 95% N_2_ and 5% CO_2_ to induce hypoxia or oxygenated with 95% O_2_ and 5% CO_2_.

Human retina samples were obtained as previously described, placed in the specimen holder, and initially perfused with oxygenated bicarbonate-buffered Ames’ media (bubbled with 95% O_2_ and 5% CO_2_) for 20 min. The perfusion solution was then switched to bicarbonate-buffered Ames’ media bubbled with 95% N_2_ and 5% CO_2_ to induce hypoxia for one hour, after which light responses were recorded. Oxygenated media was then reintroduced for 15 minutes to allow the retina to recover, after which restoration of light responses was assessed.

### ERG Recording Protocol

*Ex vivo* ERG [49] was recorded as described previously [6], [50]. Human or mouse retinas were carefully removed from the underlying RPE/choroid and mounted on the *ex vivo* ERG specimen holder, as described previously. The retinas were superfused with Ames’ media containing 100 μM BaCl_2_ to block K^+^ channels on Müller glial cell with or without 40 μM DL-AP4, which blocks signal transmission between photoreceptors and ON-bipolar cells [27], [51]. This allowed the recording of isolated photoreceptor light responses and combined photoreceptor and ON-bipolar cell responses, respectively, and the calculation of isolated ON-bipolar cell responses. We plotted photoreceptor and ON-bipolar cell amplitudes as a function of light flash strengths (photons µm^-2^) and displayed arithmetic means ± standard error of the mean.

### Small-Volume Closed Perfusion System

We built a closed perfusion *ex vivo* electroretinography ERG system that uses a peristaltic pump to circulate small volumes of perfusate. Peripheral retinal pieces from human donor eyes were used with enucleation delays of 1 (UFT8, see Table 1) to 4 (D54_2 and D55, see Table 1) hours. After overnight incubation of the eyecups, rod photoreceptor responses were recorded at multiple time points over a 12-hour period. The perfusate consisted of HEPES-buffered Ames’ media at 32.0 ± 0.5°𝐶 , an 1-micron pore filter, and Penicillin-Streptomycin (Pen-Strep purchased from Sigma-Aldrich) to prevent bacterial contamination. To isolate the photoreceptor component of the ERG response, 100 µM BaCl_2_ and 40 µM DL-AP4 were introduced. The dark-adapted retina was stimulated with 505 nm full-field 5 ms flashes. Two dripping burettes were utilized to break the electrical connection and dampen mechanical pulsations from the peristaltic pump. The perfusion system was grounded using Ag/AgCl electrodes (model EP2, World Precision Instruments) through agar gel bridges within the Faraday cage.

---

**Supplementary Figure 1.**
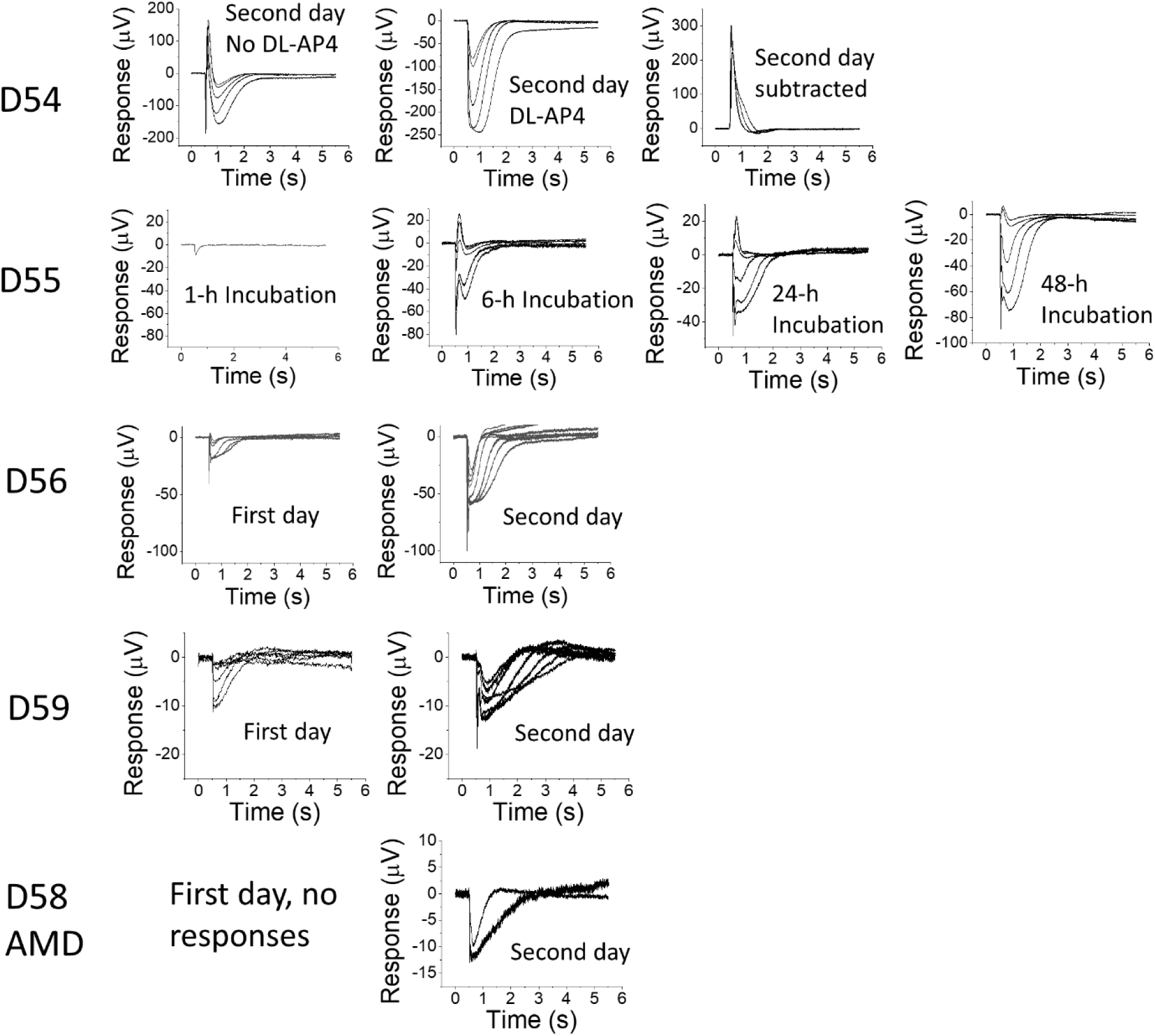
Impact of overnight eyecup incubation on the light responses in research donor eyes enucleated 3-4 hours postmortem. Light responses from peripheral retina samples from five research donors (D54, D55, D56, and D58_AMD) from whom eyes were recovered 3-4 hours postmortem (see Table 1 for donor details). D54 and D58_AMD recordings were conducted only after overnight incubation in oxygenated Ames’. Unless indicated otherwise, samples were perfused with Ames’ media containing 100 μM Ba^2+^. For D54, responses are shown in the absence (left) and presence (middle) of 40 μM DL-AP4. The rightmost panel for D54 shows responses derived by subtracting data in the middle panel from that in the left. Light flash strengths (green photons μm^-2^): D54, 19 – 1,024; D55 (6-h, 24-h and 48-h): 19 – 2,050; D59, First day: 2,300 – 2,500,000, Second day: 19 – 4,100; D56, First day: 57 – 3,900; Second day: 57 – 7,800; D58_AMD: 19 – 16,400.

**Supplementary Figure 2.**
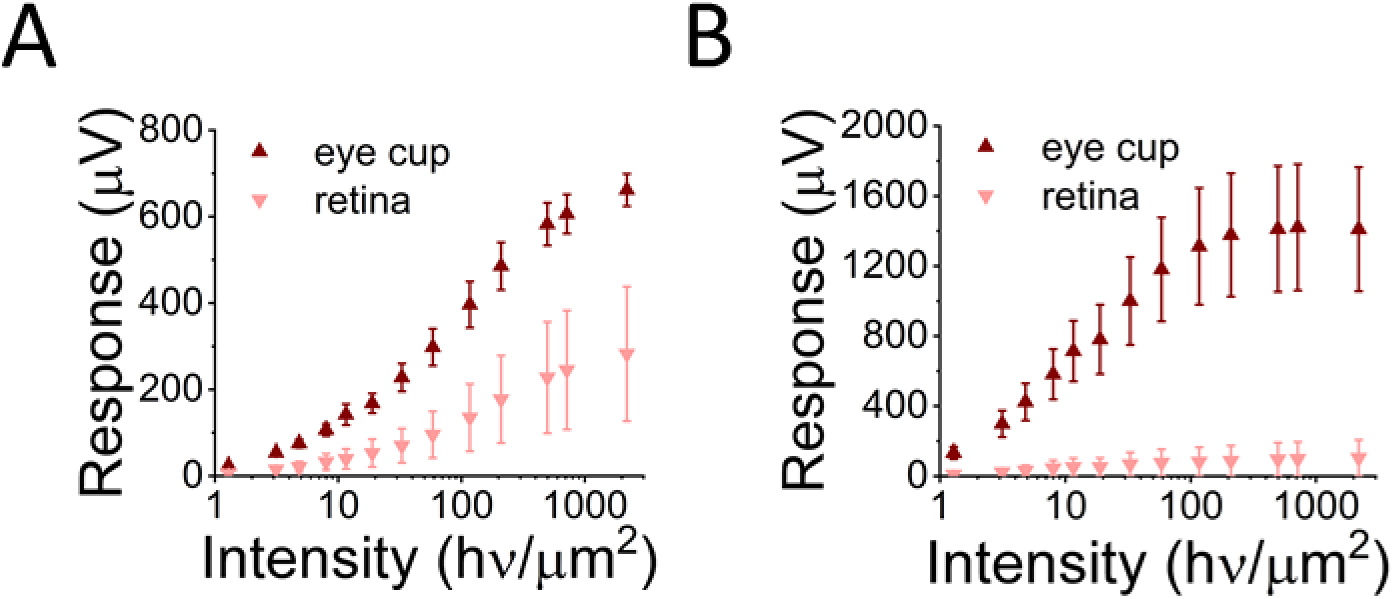
RPE dependence of retinal function after overnight storage. (a) *Ex vivo* ON-bipolar and (b) photoreceptor cell light responses after overnight storage of eye cups (n=4) or isolated retinas without underlying RPE (N=3).

**Supplementary Figure 3.**
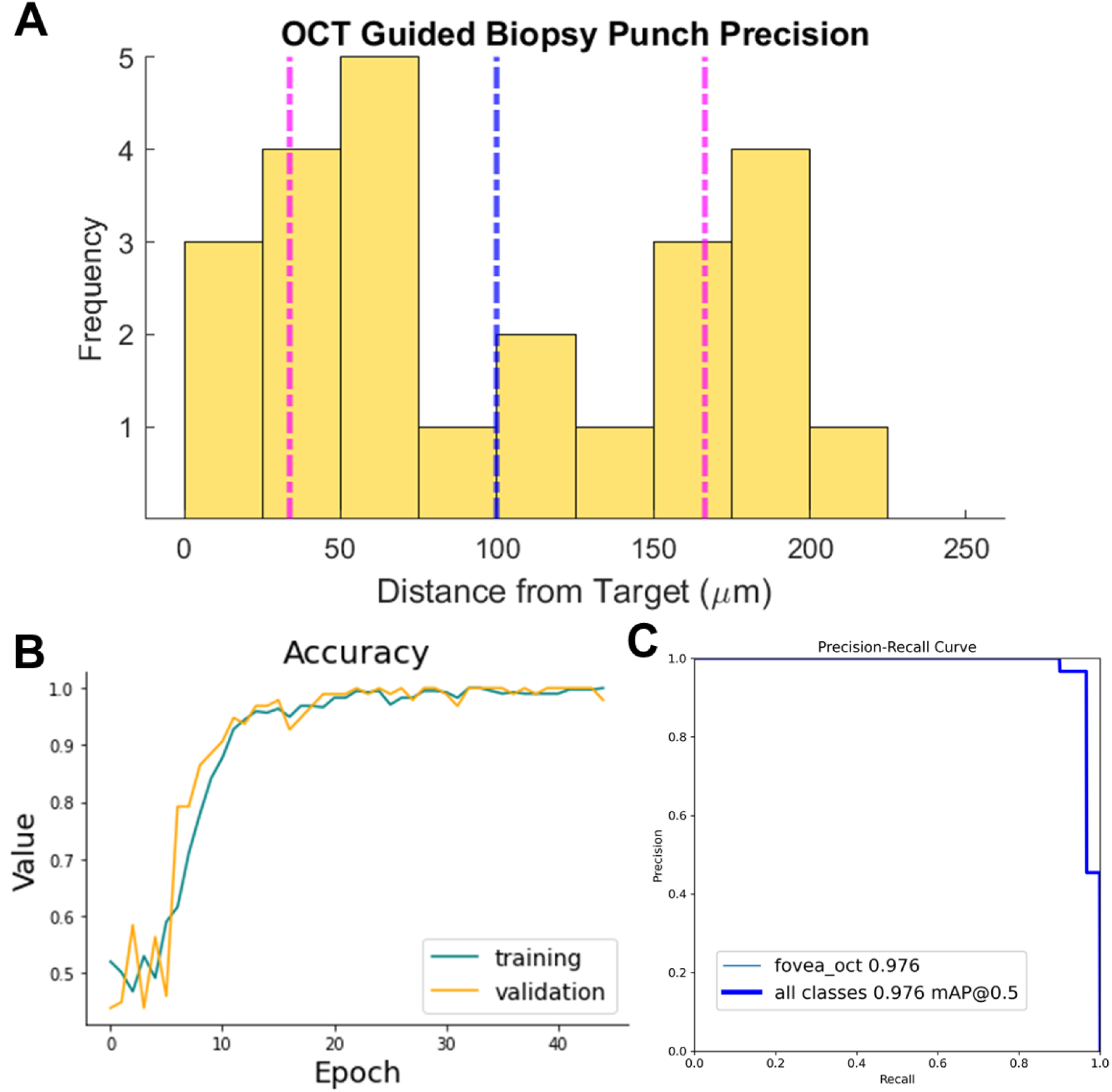
OCT and AI model accuracy and precision. (a) Bar chart of OCT precision test displaying the distances between each trial punch (N=24) from the target, with a mean (blue) of 100 ± 66.4 μm (purple). The range of each histogram bin is 25 μm. (b) Accuracy of classification model training on 592 images. The best checkpoint was used for the final model, where accuracy in training and validation was 100%. (c) Precision-Recall curve of object detection model trained on 291 images. The Mean Average Precision at 50% IoU threshold is 97.6%.

**Supplementary Figure 4.**
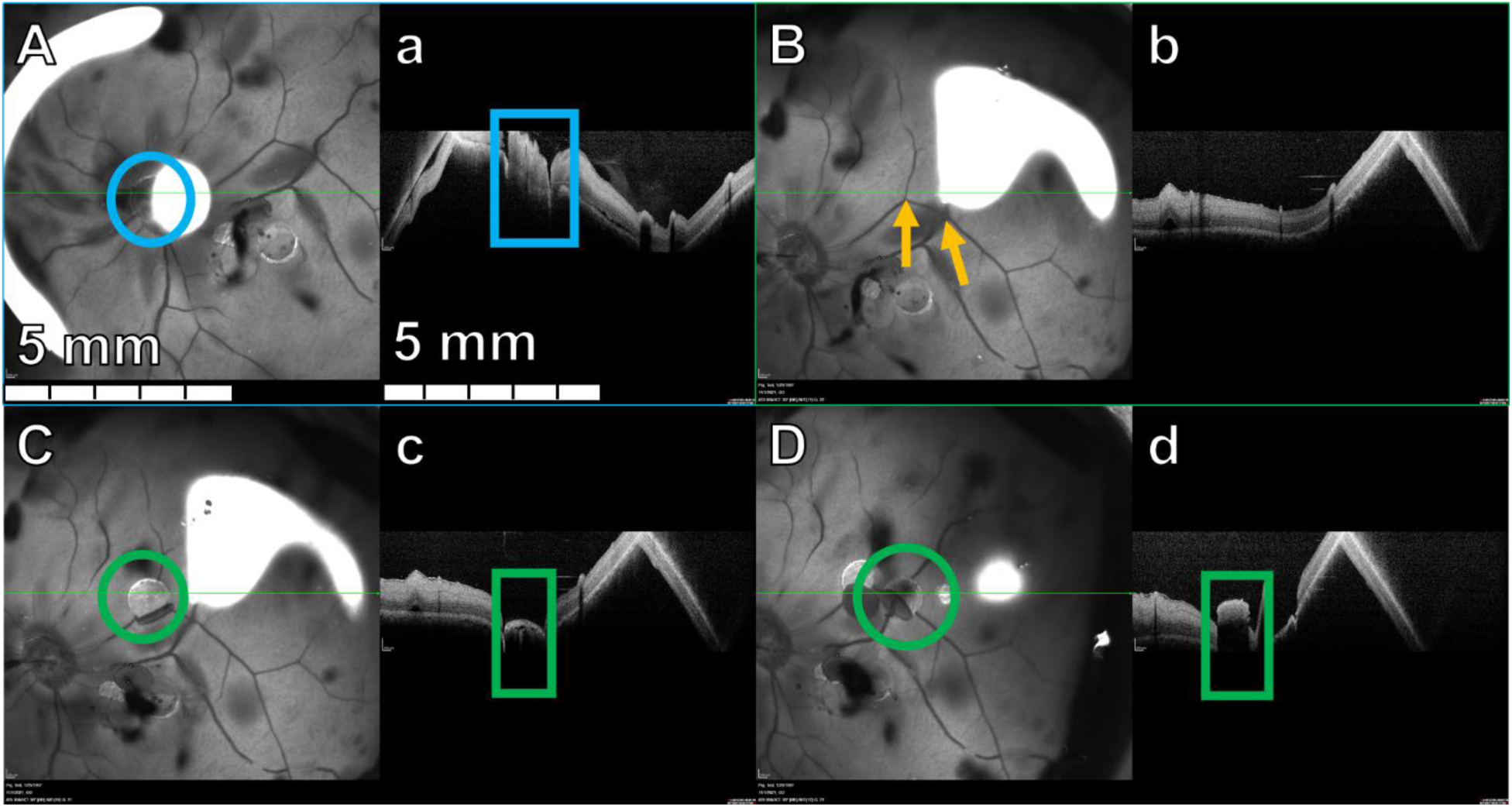
Sampling precision in porcine retina. Fundus images (capital letters) are captured with infrared imaging. OCT images (lowercase letters) show cross sectional analysis (relative to b-scan; green line). A reference punch is made (A,a; blue). Two targeted areas are selected (orange arrows in (B)). Biopsy punches are taken (green), (C,c), and (D,d). 100% of targeted areas were successfully sampled (n=9).

**Supplementary Figure 5.**
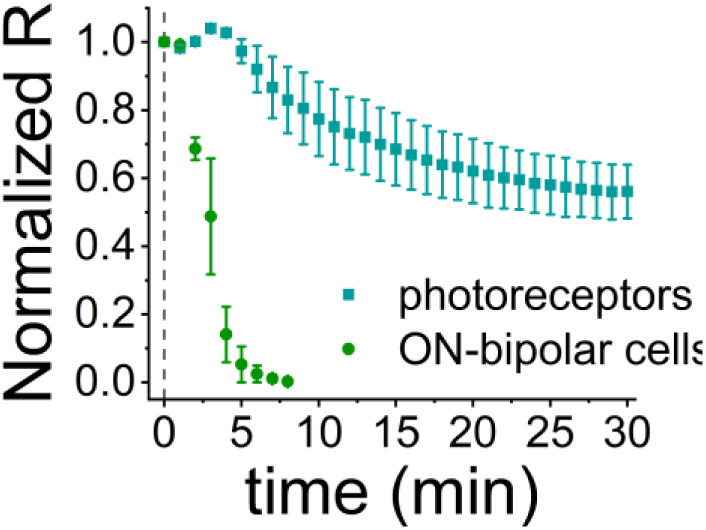
Sensitivity of retinal neurons to hypoxia. Photoreceptor (blue) and ON-bipolar cell responses (green) during hypoxia (2.5% O_2_), normalized to the response amplitudes before the onset of hypoxia (mean ± SEM, n=3 for photoreceptors, n=2 for ON-bipolar cells).

**Supplementary Figure 6.**
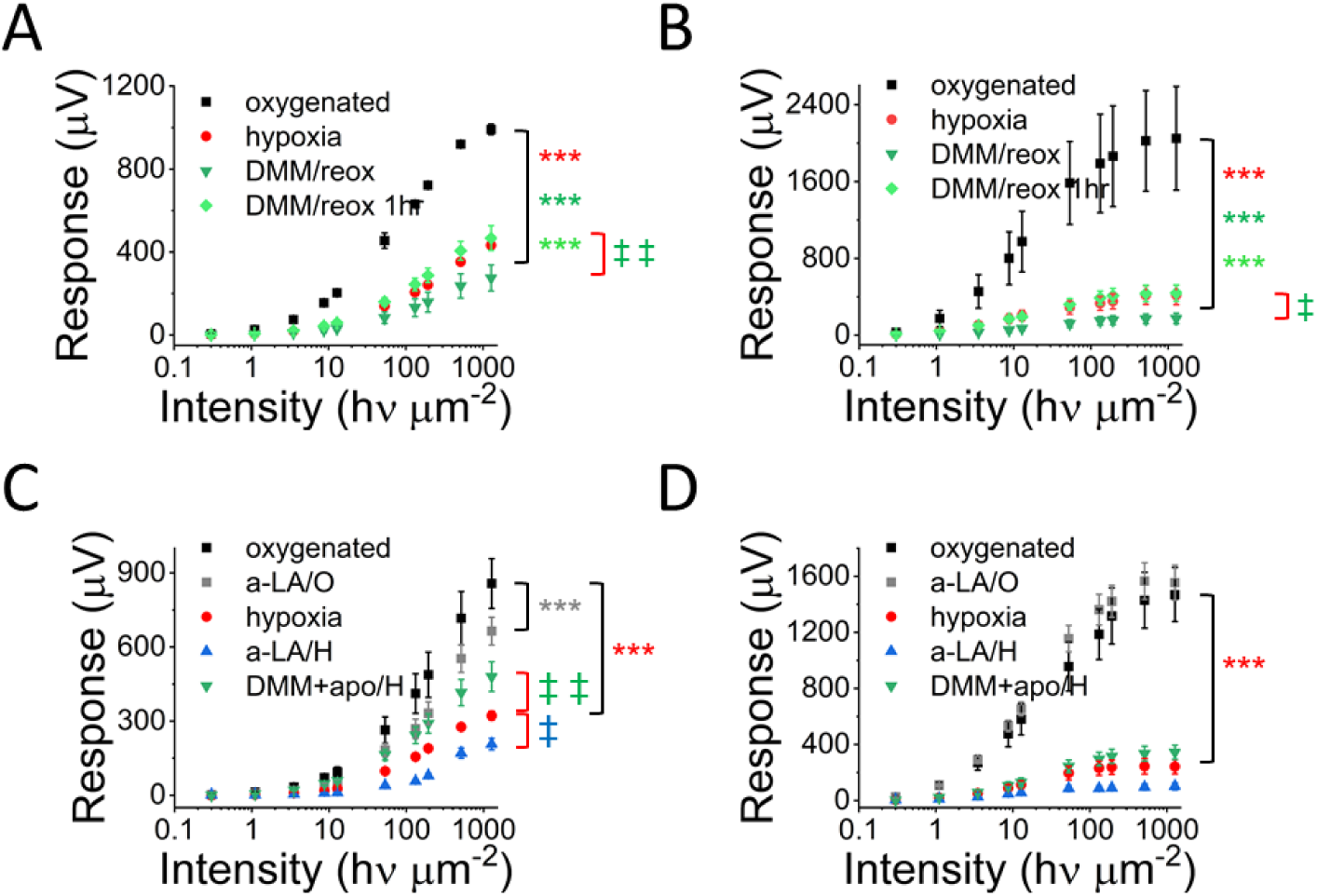
Antioxidant drugs during hypoxia and reoxygenation. Photoreceptor (A) and ON-bipolar cell light responses (B) after three hours of hypoxia with overnight incubation in oxygenated Ames’ media in the absence or presence of dimethyl malonate (DMM) for 10 min before and one hour during reoxygenation or only during overnight reoxygenation (n=2-3). Photoreceptor (C) and ON-bipolar cell light responses (D) in oxygenated media or after three hours of hypoxia with overnight incubation in oxygenated Ames’ media in the absence or presence of α-lipoic acid (a-LA) or DMM+apocynin (DMM+apo, n=4- 5).

**Supplementary Table 1.** Donor information. Preservation delay: Time from cross-clamp (organ donors) or death (research donors) until preservation in Ames’ media; SIGSW: self-inflicted gunshot wound; ICB/ICG: intracranial bleeding/hemorrhage; COPD: chronic obstructive pulmonary disease.

| Donor ID | Donor type | Preservation delay/mins | Age/sex | Cause of death | Prior medical conditions |
| --- | --- | --- | --- | --- | --- |
| UFT2 | Organ donor | 54 | 56/male | SIGSW | Type 2 diabetes. no ocular. |
| UFT3 | Organ donor | 50 | 63/female | ICB/ICH | Astigmatism. |
| UFT6 | Organ donor | 33 | 37/female | Respiratory arrest | Asthma, morbid obesity, lactic acid increased. No ocular. |
| UFT7 | Organ donor | 22 | 54/female | ICB/ICH | Cardiomyopathy. No ocular. |
| UFT8 (Closed perfusion) | Organ donor | 68 | 38/male | Anoxic brain injury | Cardiomyopathy, hypercholesterolemia. Glaucoma. |
| UFT10 | Organ donor | 77 | 57/female | ICB/ICH | Ischemic right carotid artery stroke |
| ARPAH1 | Organ donor | 46 | 61/female | Cerebrovascular accident | LASIK. |
| ARPAH2 | Organ donor | 73 | 73/female | Acute respiratory failure leading to acquired brain injury | Type 2 diabetes, intraocular lenses in both eyes. |
| D42 | Research donor | 250 | 94/male | Pneumonia | Congestive heart failure, diabetes mellitus, diabetic eye disease |
| D53 | Research donor | 180 | 68/female | Cancer | Cardiomyopathy, sepsis |
| D55 | Research donor | 247 | 89/male |  |  |
| D39 AMD | Research donor | 173 | 90/female |  | AMD |
| D46 AMD | Research donor | 346 | 75/male | Surgical complications | AMD, glaucoma |
| D48 AMD | Research donor | 208 | 89/female | Acute Cardiac Event |  |
| D51 AMD | Research donor | 161 | 96/female | Myocardial infarction | AMD, glaucoma |
| D57 AMD | Research donor | 219 | 93/female |  |  |
| D54 | Research donor | 284 | 77/female | Cancer | Abnormal EKG, hyperlipidemia, hypertension |
| D54_2 (Closed perfusion) | Research donor | 258 | 81/Male | Acute Cardiac Event | Chronic venous insufficiency, aortic stenosis, hypoxia, morbid obesity. |
| D55 (Closed perfusion) | Research donor | 249 | 89/male | Not known | Not known |
| D59 | Research donor | 231 | 78/male | Sepsis | Stroke, lactic acidosis. Vision changes, vision loss. |
| D56 | Research donor | 232 | 82/female | Ischemic bowel | Liver cirrhosis, stroke, COPD, home O2 therapy.<br>Ocular stroke, visual field loss. |
| D58_AMD | Research donor | 219 | 99/female | Pneumonia | AMD. |
| RD61 | Research donor | 259 | 88/female | Abdominal or thoracic aortic aneurysm | Eyelid OS drooping, AMD. |

## Bibliography

[1] J. Borjigin et al., “Surge of neurophysiological coherence and connectivity in the dying brain,” Proc Natl Acad Sci U S A, vol. 110, no. 35, pp. 14432–14437, Aug. 2013, doi: 10.1073/pnas.1308285110.

[2] S. L. Cole and E. Corday, “Four-minute limit for cardiac resuscitation,” J Am Med Assoc, vol. 161, no. 15, pp. 1454–1458, Aug. 1956, doi: 10.1001/jama.1956.02970150022005.

[3] N. N. Osborne, R. J. Casson, J. P. M. Wood, G. Chidlow, M. Graham, and J. Melena, “Retinal ischemia: mechanisms of damage and potential therapeutic strategies,” Prog Retin Eye Res, vol. 23, no. 1, pp. 91–147, Jan. 2004, doi: 10.1016/j.preteyeres.2003.12.001.

[4] Z. Vrselja et al., “Restoration of brain circulation and cellular functions hours post-mortem,” Nature, vol. 568, no. 7752, pp. 336–343, Apr. 2019, doi: 10.1038/s41586-019-1099-1.

[5] D. Andrijevic et al., “Cellular recovery after prolonged warm ischaemia of the whole body,” Nature, vol. 608, no. 7922, pp. 405–412, Aug. 2022, doi: 10.1038/s41586-022-05016-1.

[6] F. Abbas et al., “Revival of light signalling in the postmortem mouse and human retina,” Nature, vol. 606, no. 7913, Art. no. 7913, Jun. 2022, doi: 10.1038/s41586-022-04709-x.

[7] C. Ingensiep, K. Schaffrath, B. Denecke, P. Walter, and S. Johnen, “A multielectrode array-based hypoxia model for the analysis of electrical activity in murine retinae,” J Neurosci Res, vol. 99, no. 9, pp. 2172–2187, Sep. 2021, doi: 10.1002/jnr.24899.

[8] S. S. Hayreh, M. B. Zimmerman, A. Kimura, and A. Sanon, “Central retinal artery occlusion. Retinal survival time,” Exp Eye Res, vol. 78, no. 3, pp. 723–736, Mar. 2004, doi: 10.1016/s0014-4835(03)00214-8.

[9] K. Reinhard et al., “Hypothermia Promotes Survival of Ischemic Retinal Ganglion Cells,” Invest Ophthalmol Vis Sci, vol. 57, no. 2, pp. 658–663, Feb. 2016, doi: 10.1167/iovs.15-17751.

[10] L. S. Mure, F. Vinberg, A. Hanneken, and S. Panda, “Functional diversity of human intrinsically photosensitive retinal ganglion cells,” Science, vol. 366, no. 6470, pp. 1251–1255, Dec. 2019, doi: 10.1126/science.aaz0898.

[11] C. Chen, G. Singh, R. Madike, and S. Cugati, “Central retinal artery occlusion: a stroke of the eye,” Eye (Lond), vol. 38, no. 12, pp. 2319–2326, Aug. 2024, doi: 10.1038/s41433-024-03029-w.

[12] A. K. Rudkin, A. W. Lee, E. Aldrich, N. R. Miller, and C. S. Chen, “Clinical characteristics and outcome of current standard management of central retinal artery occlusion,” Clinical & Experimental Ophthalmology, vol. 38, no. 5, pp. 496–501, 2010, doi: 10.1111/j.1442-9071.2010.02280.x.

[13] S. Cugati, D. D. Varma, C. S. Chen, and A. W. Lee, “Treatment Options for Central Retinal Artery Occlusion,” Curr Treat Options Neurol, vol. 15, no. 1, pp. 63–77, 2013, doi: 10.1007/s11940-012-0202-9.

[14] S. S. Hayreh and M. B. Zimmerman, “Central retinal artery occlusion: visual outcome,” Am J Ophthalmol, vol. 140, no. 3, pp. 376–391, Sep. 2005, doi: 10.1016/j.ajo.2005.03.038.

[15] S. Saeid et al., “Closed-perfusion transretinal ERG setup for preclinical drug and nanostructure testing,” IEEE Trans Biomed Eng, vol. PP, Nov. 2024, doi: 10.1109/TBME.2024.3493616.

[16] L. Cangiano and S. Asteriti, “An Ex Vivo Electroretinographic Apparatus for the mL-Scale Testing of Drugs to One Day and Beyond,” Int J Mol Sci, vol. 24, no. 14, p. 11346, Jul. 2023, doi: 10.3390/ijms241411346.

[17] C. S. Cowan et al., “Cell Types of the Human Retina and Its Organoids at Single-Cell Resolution,” Cell, vol. 182, no. 6, pp. 1623–1640.e34, Sep. 2020, doi: 10.1016/j.cell.2020.08.013.

[18] X. Zhong et al., “Generation of three-dimensional retinal tissue with functional photoreceptors from human iPSCs,” Nat Commun, vol. 5, p. 4047, Jun. 2014, doi: 10.1038/ncomms5047.

[19] K. C. Eldred et al., “Thyroid hormone signaling specifies cone subtypes in human retinal organoids,” Science, vol. 362, no. 6411, p. eaau6348, Oct. 2018, doi: 10.1126/science.aau6348.

[20] Y. C. Leong, V. Di Foggia, H. Pramod, M. Bitner-Glindzicz, A. Patel, and J. C. Sowden, “Molecular pathology of Usher 1B patient-derived retinal organoids at single cell resolution,” Stem Cell Reports, vol. 17, no. 11, pp. 2421–2437, Nov. 2022, doi: 10.1016/j.stemcr.2022.09.006.

[21] V. Busskamp et al., “Genetic reactivation of cone photoreceptors restores visual responses in retinitis pigmentosa,” Science, vol. 329, no. 5990, pp. 413–417, Jul. 2010, doi: 10.1126/science.1190897.

[22] A. Saha, E. Capowski, M. A. Fernandez Zepeda, E. C. Nelson, D. M. Gamm, and R. Sinha, “Cone photoreceptors in human stem cell-derived retinal organoids demonstrate intrinsic light responses that mimic those of primate fovea,” Cell Stem Cell, vol. 29, no. 3, pp. 460–471.e3, Mar. 2022, doi: 10.1016/j.stem.2022.01.002.

[23] H. Kolb, R. F. Nelson, P. K. Ahnelt, I. Ortuño-Lizarán, and N. Cuenca, “The Architecture of the Human Fovea,” in *Webvision: The Organization of the Retina and Visual System*, H. Kolb, E. Fernandez, and R. Nelson, Eds., Salt Lake City (UT): University of Utah Health Sciences Center, 1995. Accessed: Jun. 03, 2024. [Online] . Available: http://www.ncbi.nlm.nih.gov/books/NBK554706/

[24] S. Becker, Z. L’Ecuyer, B. W. Jones, M. A. Zouache, F. S. McDonnell, and F. Vinberg, “Modeling complex age-related eye disease,” Prog Retin Eye Res, vol. 100, p. 101247, May 2024, doi: 10.1016/j.preteyeres.2024.101247.

[25] P. Witkovsky, F. E. Dudek, and H. Ripps, “Slow PIII component of the carp electroretinogram,” J Gen Physiol, vol. 65, no. 2, pp. 119–134, Feb. 1975, doi: 10.1085/jgp.65.2.119.

[26] D. A. Bolnick, A. E. Walter, and A. J. Sillman, “Barium suppresses slow PIII in perfused bullfrog retina,” Vision Res, vol. 19, no. 10, pp. 1117–1119, 1979, doi: 10.1016/0042-6989(79)90006-3.

[27] S. Nymark, H. Heikkinen, C. Haldin, K. Donner, and A. Koskelainen, “Light responses and light adaptation in rat retinal rods at different temperatures,” J Physiol, vol. 567, no. Pt 3, pp. 923–938, Sep. 2005, doi: 10.1113/jphysiol.2005.090662.

[28] P. Kofuji, P. Ceelen, K. R. Zahs, L. W. Surbeck, H. A. Lester, and E. A. Newman, “Genetic inactivation of an inwardly rectifying potassium channel (Kir4.1 subunit) in mice: phenotypic impact in retina,” J Neurosci, vol. 20, no. 15, pp. 5733–5740, Aug. 2000, doi: 10.1523/JNEUROSCI.20-15-05733.2000.

[29] J. D. Sokolowski et al., “Preclinical models of middle cerebral artery occlusion: new imaging approaches to a classic technique,” Front Neurol, vol. 14, p. 1170675, 2023, doi: 10.3389/fneur.2023.1170675.

[30] M. Renner et al., “Optic Nerve Degeneration after Retinal Ischemia/Reperfusion in a Rodent Model,” Front Cell Neurosci, vol. 11, p. 254, 2017, doi: 10.3389/fncel.2017.00254.

[31] N. Vestergaard, L. J. Cehofski, B. Honoré, K. Aasbjerg, and H. Vorum, “Animal Models Used to Simulate Retinal Artery Occlusion: A Comprehensive Review,” Transl Vis Sci Technol, vol. 8, no. 4, p. 23, Jul. 2019, doi: 10.1167/tvst.8.4.23.

[32] H. M. Kim, K. H. Park, and S. J. Woo, “Correlation of electroretinography components with visual function and prognosis of central retinal artery occlusion,” Sci Rep, vol. 10, no. 1, p. 12146, Jul. 2020, doi: 10.1038/s41598-020-68957-5.

[33] A. R. Fernández, R. Sánchez-Tarjuelo, P. Cravedi, J. Ochando, and M. López-Hoyos, “Review: Ischemia Reperfusion Injury-A Translational Perspective in Organ Transplantation,” Int J Mol Sci, vol. 21, no. 22, p. 8549, Nov. 2020, doi: 10.3390/ijms21228549.

[34] S.-B. Ong, P. Samangouei, S. B. Kalkhoran, and D. J. Hausenloy, “The mitochondrial permeability transition pore and its role in myocardial ischemia reperfusion injury,” J Mol Cell Cardiol, vol. 78, pp. 23–34, Jan. 2015, doi: 10.1016/j.yjmcc.2014.11.005.

[35] J. L. Martin et al., “Succinate accumulation drives ischaemia-reperfusion injury during organ transplantation,” Nat Metab, vol. 1, pp. 966–974, Sep. 2019, doi: 10.1038/s42255-019-0115-y.

[36] E. T. Chouchani et al., “Ischaemic accumulation of succinate controls reperfusion injury through mitochondrial ROS,” Nature, vol. 515, no. 7527, pp. 431–435, Nov. 2014, doi: 10.1038/nature13909.

[37] J. Stefanska and R. Pawliczak, “Apocynin: Molecular Aptitudes,” Mediators of Inflammation, vol. 2008, p. 106507, Dec. 2008, doi: 10.1155/2008/106507.

[38] D. Sun, W. Gao, H. Hu, and S. Zhou, “Why 90% of clinical drug development fails and how to improve it?,” Acta Pharm Sin B, vol. 12, no. 7, pp. 3049–3062, Jul. 2022, doi: 10.1016/j.apsb.2022.02.002.

[39] A. Loewa, J. J. Feng, and S. Hedtrich, “Human disease models in drug development,” Nat Rev Bioeng, vol. 1, no. 8, Art. no. 8, Aug. 2023, doi: 10.1038/s44222-023-00063-3.

[40] A. Akhtar, S. M. Gupta, S. Dwivedi, D. Kumar, Mohd. F. Shaikh, and A. Negi, “Preclinical Models for Alzheimer’s Disease: Past, Present, and Future Approaches,” ACS Omega, vol. 7, no. 51, pp. 47504–47517, Dec. 2022, doi: 10.1021/acsomega.2c05609.

[41] T. M. Dawson, T. E. Golde, and C. L. Tourenne, “Animal Models of Neurodegenerative Diseases,” Nat Neurosci, vol. 21, no. 10, pp. 1370–1379, Oct. 2018, doi: 10.1038/s41593-018-0236-8.

[42] S. Schnichels et al., “Retina in a dish: Cell cultures, retinal explants and animal models for common diseases of the retina,” Progress in Retinal and Eye Research, vol. 81, p. 100880, Mar. 2021, doi: 10.1016/j.preteyeres.2020.100880.

[43] J. Rivera and L. Tessarollo, “Genetic background and the dilemma of translating mouse studies to humans,” Immunity, vol. 28, no. 1, pp. 1–4, Jan. 2008, doi: 10.1016/j.immuni.2007.12.008.

[44] D. Ail et al., “Inducible nonhuman primate models of retinal degeneration for testing end-stage therapies,” Sci Adv, vol. 9, no. 31, p. eadg8163, Aug. 2023, doi: 10.1126/sciadv.adg8163.

[45] D. J. Ceradini et al., “Combined Whole Eye and Face Transplant: Microsurgical Strategy and 1-Year Clinical Course,” JAMA, vol. 332, no. 18, pp. 1551–1558, Nov. 2024, doi: 10.1001/jama.2024.12601.

[46] “NYU Langone Health Performs World’s First Whole-Eye & Partial-Face Transplant,” NYU Langone News. Accessed: May 12, 2024. [Online] . Available: https://nyulangone.org/news/nyu-langone-health-performs-worlds-first-whole-eye-partial-face-transplant

[47] “THEA | ARPA-H.” Accessed: May 25, 2024. [Online]. Available: https://arpa-h.gov/research-and-funding/programs/thea

[48] R. N. Frank and J. E. Dowling, “Rhodopsin photoproducts: effects on electroretinogram sensitivity in isolated perfused rat retina,” Science, vol. 161, no. 3840, pp. 487–489, Aug. 1968, doi: 10.1126/science.161.3840.487.

[49] F. Abbas, F. Vinberg, and S. Becker, “Optimizing the Setup and Conditions for Ex Vivo Electroretinogram to Study Retina Function in Small and Large Eyes,” J Vis Exp, no. 184, Jun. 2022, doi: 10.3791/62763.

[50] F. Vinberg and V. Kefalov, “Simultaneous ex vivo functional testing of two retinas by in vivo electroretinogram system,” J Vis Exp, no. 99, p. e52855, May 2015, doi: 10.3791/52855.

